# Human sialidase activity is vital for dengue virus serotype 2 infection

**DOI:** 10.1101/2022.10.05.511017

**Authors:** Laura A. St Clair, Padmasri G. Pujari, Rushika Perera

## Abstract

The human sialidase enzymes (or neuraminidases, NEU1-4) are glycoside hydrolases that catalyze the removal of sialic acid residues from glycoconjugates, including many bioactive glycoproteins and glycolipids. Through their physiochemical effect on glycoconjugates, sialic acid residues are thought to play vital roles in the control of cellular signaling. In previous studies, it was demonstrated that NEU1-4 activity was increased in cells infected with dengue virus serotype 2 (DENV2). Additionally, it was demonstrated that the DENV2 NS1 protein was sufficient for inducing increased NEU1-4 activity in both *in vivo* and *in vitro* models, and that this increased activity was linked to endothelial hyperpermeability and vascular leakage, a hallmark of severe dengue disease. However, the role of increased NEU1-4 activity in the viral lifecycle was not understood. Here, we used siRNA-mediated loss of function studies to evaluate the effect of inhibition of sialidase activity on the DENV2 lifecycle. Our analyses uncovered that apart from their importance for viral pathogenesis, NEU1-4 activity was vital for DENV2 viral replication and egress. Moreover, we characterized the inter-relationship between NEU 1-4, and determined that there was a transcriptional dependency of NEU1-3 on NEU4.

## Introduction

Sialic acids (SIAs) are a family of over 50 structurally distinct, sugar molecules with a 9-carbon backbone that are most often found at the terminal position of glycolipids and glycoproteins [1–4]. Given their ubiquitous occurrence, especially in cell surface molecules, SIAs are some of the most versatile and influential regulatory molecules in cell biology [1–4]. SIAs function either as ligands, masking molecules, or as physicochemical effectors of both their attached glycoconjugates and surrounding molecules due to their strong negative charge [1–4]. As such, SIAs are vital for cell signaling, cellular recognition, cell-to-cell communication, immune cell signaling and modulation, cell differentiation, cellular trafficking, and structural conformation of glycoconjugates [1–4]. As the roles of SIAs in biological functions are dynamic, so must be the balance between sialylated and desialylated states of glycoconjugates. This balance is mediated by sialytransferases and sialidases (neuraminidases), respectively [1–4]. In humans, there are 4 known sialidases (NEU1-4). Interestingly, we and others have shown that the DENV2 nonstructural protein 1 (NS1) is necessary and sufficient to cause increased activity of the human sialidase enzymes (neuraminidase 1-4, NEU1-4) in both *in vitro* and *in vivo* models [9,10]. Moreover, the upregulated sialidase activity was associated with increased endothelial hyperpermeability in *in vitro* models, and vascular leakage coupled with increased morbidity and mortality in an *in vivo* mouse model [9,10]. These studies revealed a link between increased sialidase activity and DENV pathogenesis; however, it remains unclear what advantage increased sialidase activity plays in the virus lifecycle.

In this study, we sought to characterize the role of the four human sialidase enzymes, NEU1-4, on the DENV2 lifecycle in human hepatoma cells (Huh7s). We used siRNA-mediated loss of function analysis to systematically examine the role each enzyme plays during the viral replication cycle. We uncovered that these enzymes do not only contribute to viral pathogenesis, but also play a critical role in viral replication and egress. Importantly, this study highlighted yet uncovered roles of sialic acids and sialidase enzyme activity in the DENV2 lifecycle.

## Results

### Sialidase activity is vital for DENV2 replication and viral release

Neuraminidases were previously shown to be upregulated by DENV2 NS1 protein [9,10]. This upregulation contributed to the breakdown of endothelial barrier integrity that leads to the vascular leakage observed in severe dengue disease [9,10]. However, it was unknown whether this upregulation contributed only to viral pathogenesis or also played a role in the DENV2 lifecycle. Moreover, these previous results were intriguing given that NEU1-4 are found in different subcellular compartments (Figure 1A), and have different substrate specificities (Table 1) [11,12, reviewed in 13–15]. We hypothesized that NEU1-4 may each have specific, independent roles within the viral lifecycle. Thus, we first determined the effect of siRNA-mediated loss of function of NEU1-4 on DENV2 release from Huh7 cells. Interestingly, we found that inhibition of each enzyme resulted in a significant decrease (1-2 log reduction) in infectious virus release from cells (Figure 1B). Similarly, we also saw a significant decrease in DENV2 genome replication (Figure 1C), however, not to the same magnitude as viral release. Knockdown (KD) of NEU1-4 was validated by measuring mRNA expression using qRT-PCR (Supplemental Figure 1A-B).

**Table 1.**
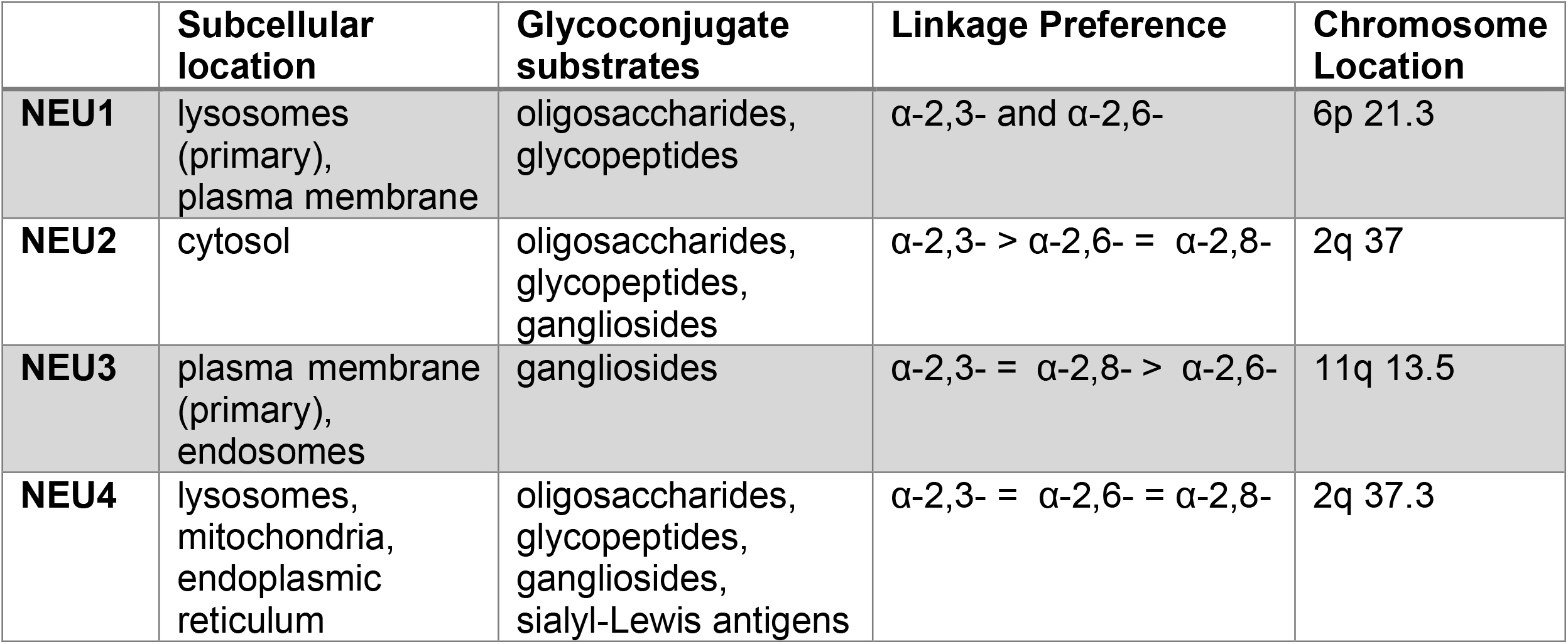
Properties of Human Sialidases.

**Figure 1.**
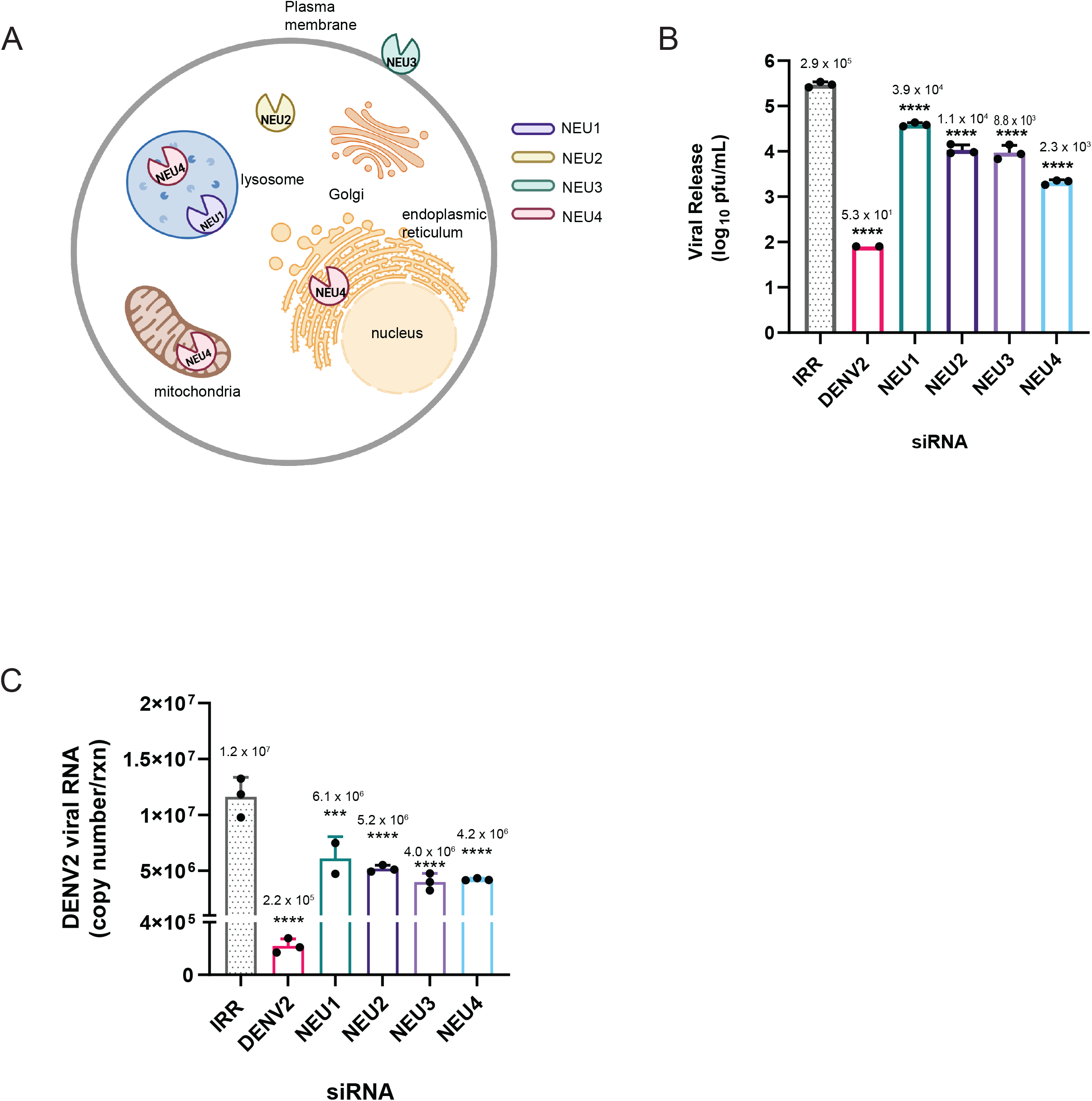
Subcellular Location of human sialidases and preliminary loss of function analysis of NEU1-4. (A) Primary subcellular location of human sialidase enzymes. B-C: Huh7 cells were transfected with pooled siRNAs targeting NEU1-4, a DENV2-specific positive control siRNA, and an irrelevant (non-targeting) negative control siRNA, followed by DENV2 infection (MOI = 0.3) for 24 hr. (B) Supernatants were titrated via plaque assay on BHK-21 cells to measure infectious virus release. (C) Cells were collected and copy number of viral RNA was measured via qRT-PCR and normalized to the RPLP0 housekeeping gene. NEU1: human neuraminidase 1, NEU2: human neuraminidase 2, NEU3: neuraminidase 3, NEU4: neuraminidase 4, IRR: irrelevant control, DENV2: dengue virus serotype 2. (A: image generated using Biorender.com, B-C: one-way ANOVA with Dunnett’s multiple comparison’s test: * = p≤0.05, ** = p≤0.01, ***= p≤0.001, **** = p≤0.0001.)

### Viral egress is attenuated by downregulation of NEU1-4

We next sought to determine whether the reduction of viral genome replication and infectious virus released from NEU1-4 KD samples was due to defects in viral egress or infectivity. To evaluate the effect of NEU1-4 KD on particle infectivity, we measured the ratio of total particles released (genome equivalents) to infectious particles released (viral titer) in the supernatant as previously described [6]. An increase in the particle/pfu ratio would signify impaired particle maturation as mature virus particles are more infectious than immature virus particles. We determined that loss of function of NEU1-4 had no impact on the infectivity of virus particles released from NEU1-4 KD cells (Figure 2A).

**Figure 2.**
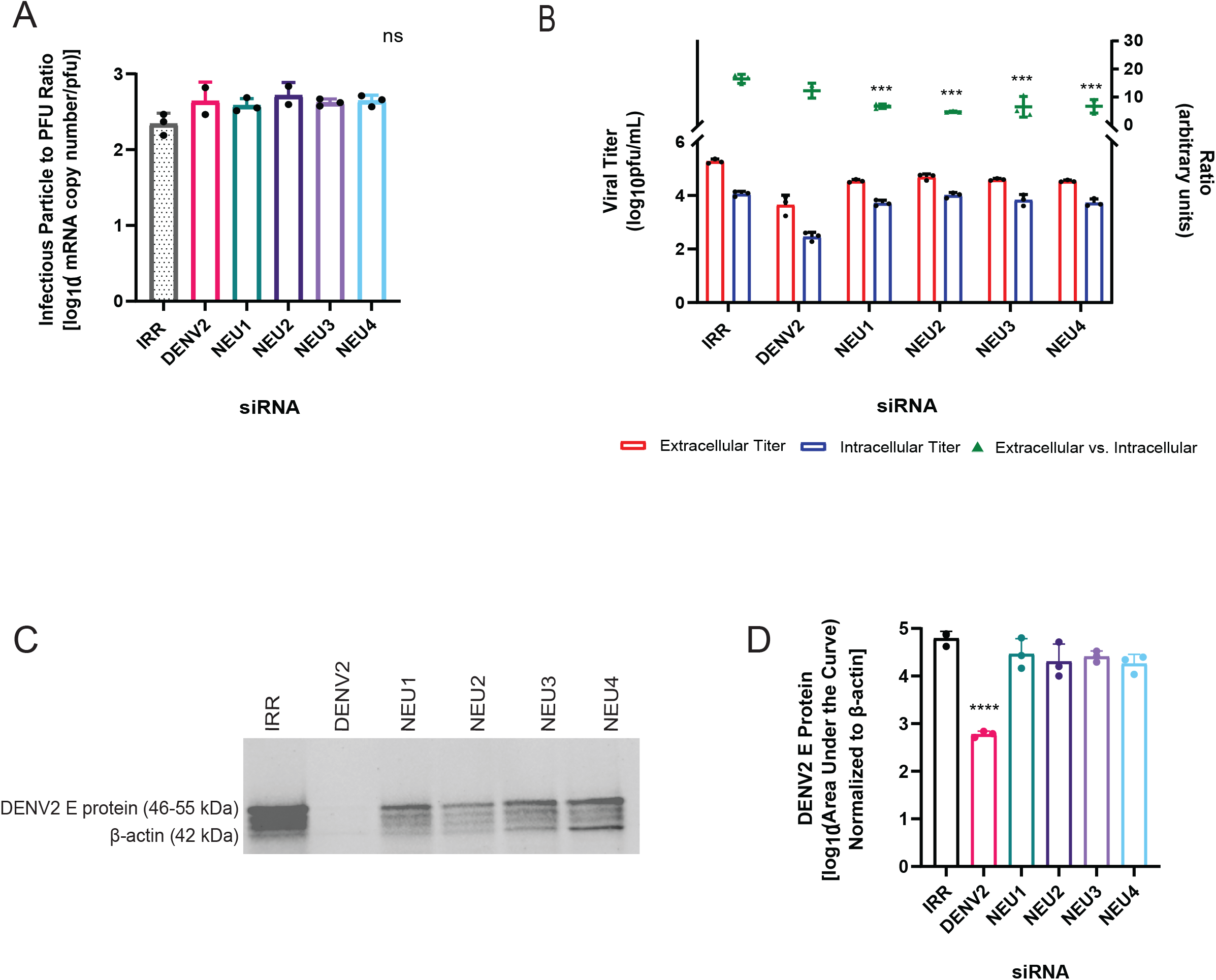
Dengue release, but not particle infective is affected by loss of function of NEU1-4. Huh7 cells were transfected with siRNAs targeting NEU1-4 and indicated controls, followed by infection with DENV2 (24 hr, MOI = 0.3). (A) At 24 hpi, viral supernatants were collected and split to be analyzed via plaque assay for infectious virus released from cells, and via qRT-PCR to measure total viral particles release (genome equivalents) release. A ratio of infectious virus particle to viral titer was taken to determine the effect of NEU1-4 KD on DENV2 particle infectivity. (B) At 24 hpi, viral supernatants were collected, and then cells were trypsinized, collected and counted. The cells were subjected to freeze-thaw lysis, and then both the viral supernatants and cell fractions were analyzed via plaque assay to examine the effect of NEU1-4 KD on viral release. A ratio of extracellular vs. intracellular virus was calculated and is displayed on the right y-axis. Extracellular and intracellular titer values were normalized to total cell count. (C) At 24 hpi, cells were collected and the amount of DENV2 E protein in each sample was measured via Western Blot. (D) Western blot was analyzed using image under the curve analysis in ImageJ, and normalized to β-actin to determine the relative amount of DENV2 E protein in each sample. (A, B, D: one-way ANOVA with Dunnett’s multiple comparison’s test: * = p≤0.05, ** = p≤0.01, ***= p≤0.001, **** = p≤0.0001.)

As neuraminidase activity is indispensable for viral egress of many viruses [16, reviewed in 17,18], we next investigated whether downregulation of NEU1-4 would also inhibit DENV2 egress. Following siRNA treatment and DENV2 infection of Huh7 cells, we collected both the viral supernatant and cells (24 hpi). Cells were gently lysed using the freeze-thaw lysis method to maintain integrity of any virions not released from the cell. We then titrated both the supernatant and cell samples on BHK-21 cells and compared the amounts of infectious DENV2 in each fraction. Overall, we found a higher amount of infectious virus in the supernatant in both the IRR control and NEU1-4 KD samples compared to their corresponding internal viral titers (Figure 2B). However, the NEU1-4 KD samples did show an overall reduction in released infectious virus, confirming our previous results (Figure 1B). Intriguingly, we noticed that the internal viral titer was similar between our IRR control and NEU1-4 KD samples. Essentially, there was a significant decrease in the ratio of extracellular versus intracellular infectious DENV2 (Figure 2B). This suggested that DENV2 release may be attenuated by loss of neuraminidase activity.

To further evaluate this observation, we performed western blot analysis of DENV2-infected NEU1-4 KD cells to determine whether virus particles accumulated in cells with reduced sialidase activity. In support of our observations of similar levels of intracellular viral titer between our control and KD samples, we also observed similar levels of DENV2 E protein between our samples (Figure 2C-D). Importantly, measurement of ‘intracellular infectious titer’ and our western blot analysis of DENV2 E protein accumulation could not distinguish between infectious virus trafficking from sites of assembly in the ER to the Golgi versus infectious virus egressing from the plasma membrane following trafficking through the secretory pathway. Thus, we next performed two groups of immunofluorescence assays to clarify these results. In the first group (Figure 3A), cells were fixed in 4% paraformaldehyde (PFA), but not permeabilized to evaluate whether the accumulation viral particles was occurring on the external surface of the plasma membrane. Interestingly, we observed similar distribution of E protein at the plasma membrane in the NEU1, NEU3 and NEU4 KD samples as in the IRR control; however, in the NEU2 KD samples a significant increase in the amount of plasma membrane-associated E protein was observed. In the second group (Figure 3B), cells were fixed in 4% PFA and permeabilized to examine whether virus was accumulating inside treated cells. As expected, the IRR control had a high number of infected cells (as indicated by significant expression of DENV2 E protein). A decrease in the overall abundance of infected cells was noted in the NEU1-4 knockdown samples in support of our initial findings that NEU1-4 decreased infectious virus release and genome replication (Figure 1B-C). Interestingly, we observed that in individual infected cells in the treated samples there was a similar expression level of DENV2 E protein as the IRR control, however spread of virus to neighboring cells was not observed. We confirmed that the accumulation of E protein was occurring in cells where NEU1-4 KD was evident (Supplemental Figure 2). Taken together, these data suggest that loss of function of sialidase activity reduced viral replication and release by reducing viral egress from infected cells, thus limiting spread.

**Figure 3.**
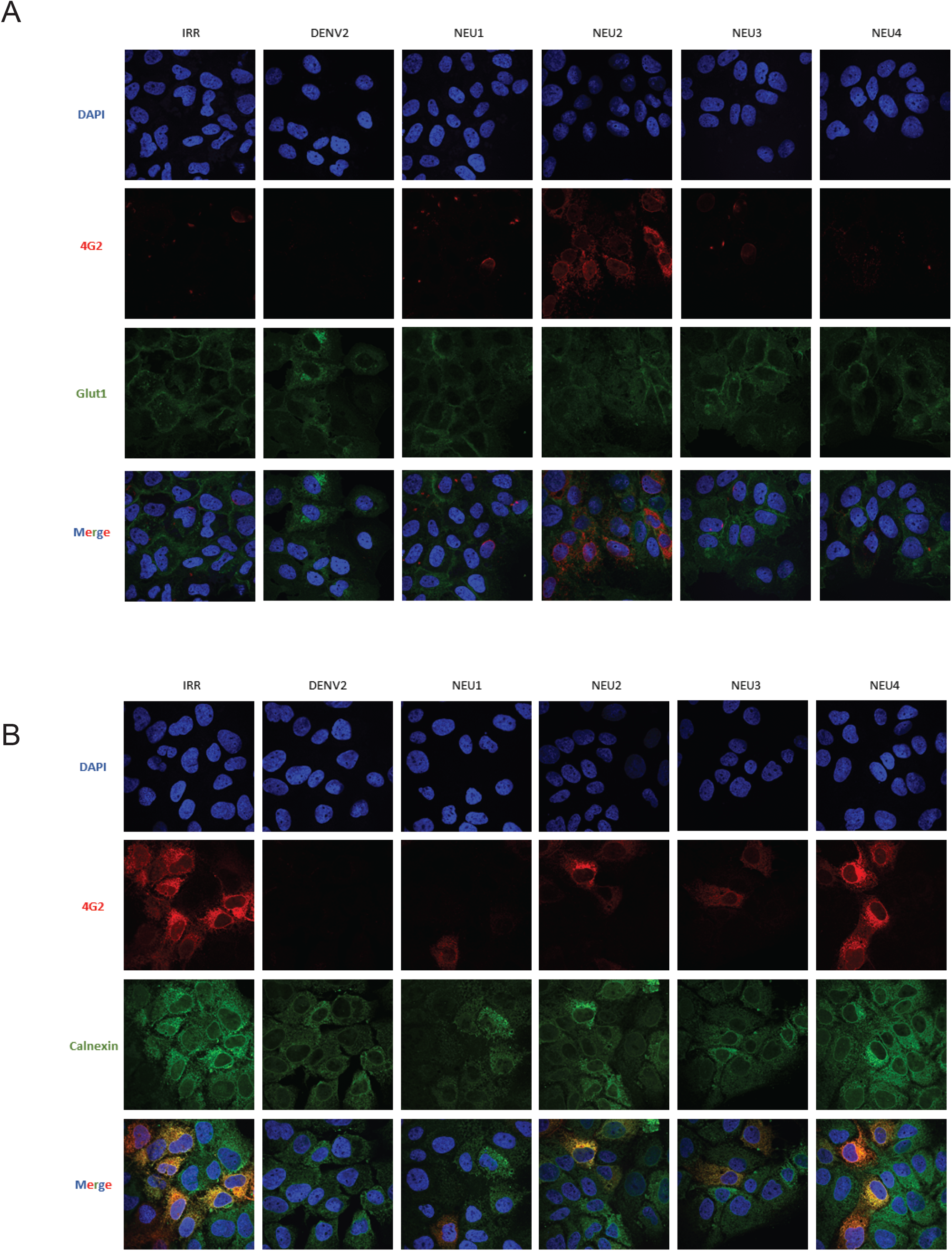
Inhibition of sialidase activity reduces DENV2 spread. (A-B) Huh7 cells were transfected with siRNAs targeting NEU1-4 and indicated controls, followed by infection with DENV2 (MOI = 0.3). At 24 hpi, cells were fixed in 4% paraformaldehyde, and either not permeabilized (A) or permeabilized (B) to determine whether NEU1-4 KD inhibits viral egress from cells. Primary antibodies used were mouse anti-4G2 (DENV2 E protein), rabbit anti-Glut1 (plasma membrane marker), or rabbit anti-Calnexin (ER marker). Cells were counterstained with goat anti-mouse AlexaFluor 488, goat anti-rabbit AlexaFluor 647, and DAPI nuclear stain and imaged on an Olympus inverted laser scanning confocal microscope and processed using Volocity software v6.3.

### NEU1-4 are transcriptionally regulated during DENV2 infection

Previously, DENV NS1 was shown to increase expression of NEU1-3 proteins by 3 hours post treatment (hpt) with recombinant NS1 [9]. This lead to a decrease in plasma membrane sialic acid residues at early timepoints (6 and 12 hpt) with a return to normal levels by 24 hpt [9]. As this previous work used recombinant NS1 protein alone to examine its effect on NEU proteins, we sought to characterize whether NEU1-4 might be transcriptionally altered during DENV2 infection. Huh7 cells were either mock- or DENV2-infected, and then collected at timepoints representing early (0, 6, 12), peak (18, 24, 36), and late (48) viral replication. We then analyzed the mRNA expression of NEU1-4 and hexokinase II at each timepoint using qRT-PCR (Figure 4). Hexokinase II was used as a positive control as it has been previously established that it is upregulated during DENV infection [19]. NEU1 mRNA expression remained at basal levels at early timepoints but began trending downward at 24 hpi in DENV2-infected cells compared to mock-infected cells. NEU2 mRNA expression remained at basal levels except at 24 hpi where a significant decrease in NEU2 expression was observed in DENV2-infected cells. These results suggested that NEU1 and NEU2 functionality may support early events of the DENV lifecycle. Interestingly, a temporal elevation in NEU3 mRNA expression every 18 hours (0, 18, and 36 hpi) was observed in DENV2 infected cells. In addition, NEU3 expression remained elevated in DENV2-infected cells compared to mock during peak and late stage. Combined, these data suggested that elevations in NEU3 expression was coincident with the start of DENV2’s replication cycle and may be critical for steps involved in viral attachment/entry. Surprisingly, NEU4 mRNA expression began to decrease at 12 hpi, and was dramatically reduced by 48 hpi. As our previous data showed that loss of function of NEU4 resulted in the most significant decrease in viral release (Figure 1B) and viral genome replication (Figure 1C), these results indicated that NEU4 must play a vital role during early events of the DENV lifecycle but may be detrimental during later timepoints.

**Figure 4.**
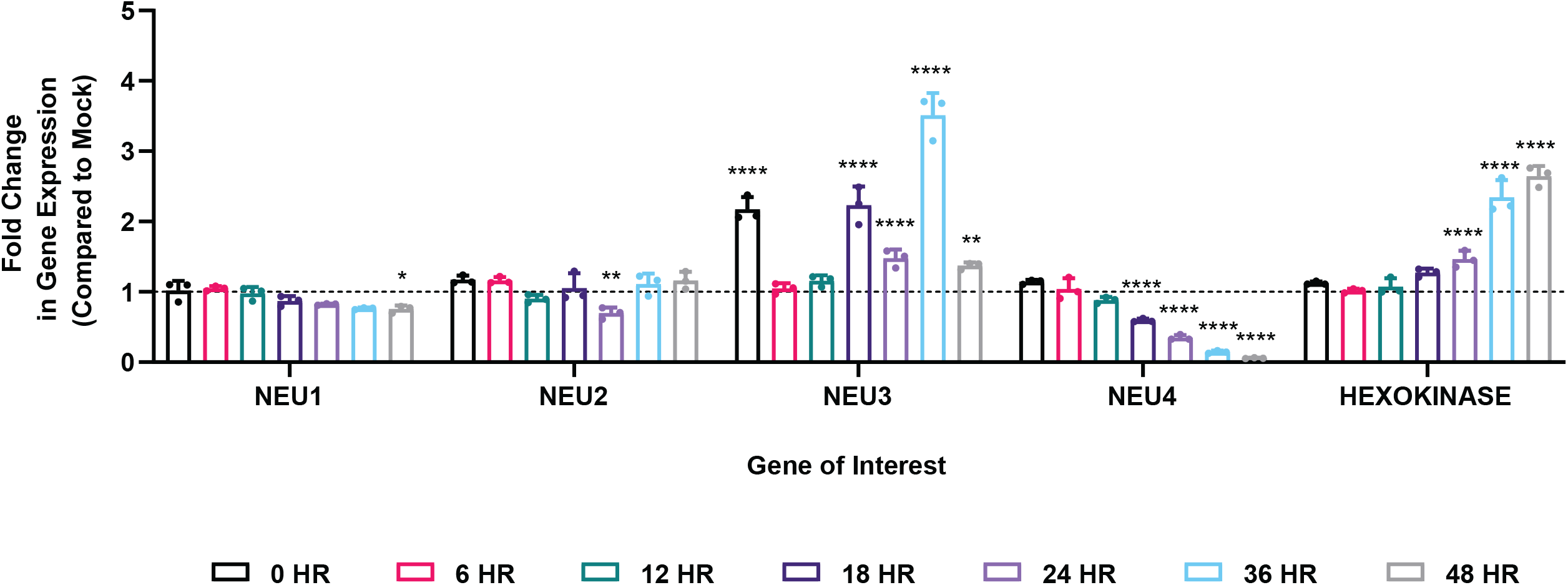
NEU1-4 mRNA is transcriptionally regulated during DENV2 infection. Huh7 cells were either mock-infected or DENV2-infected (MOI = 10), and infected cells were collected at the indicated timepoints, processed, and analyzed via qRT-PCR for mRNA expression of NEU1-4 and hexokinase II (positive control) to determine relative mRNA expression. Results were normalized to the RPLP0 housekeeping gene, and infected samples were compared to mock at each timepoint. (One-way ANOVA with Dunnett’s multiple comparison’s test: * = p≤0.05, ** = p≤0.01, ***= p≤0.001, **** = p≤0.0001.)

### Loss of function of NEU1-4 differentially affects other NEUs

NEU1-4 have been shown to translocate between cellular compartments during infection with DENVs and other viruses [9,12,20,21]. Because there is some overlap in functionality between NEU1-4 [12,14,15], we next evaluated whether each neuraminidase could compensate for loss of function of the others. Upon examining mRNA expression in our mock infected samples (Figure 5A), NEU4 expression (p < 0.01) was elevated upon knockdown of NEU1. As NEU1 and NEU4 are both found within the lysosome [22], this indicated that NEU4 may be able to compensate for loss of function of NEU1. Interestingly, in many of our samples we observed that loss of function of one NEU resulted in decreased mRNA expression of the others. In the NEU1 knockdown samples, NEU3 mRNA expression was significantly (p < 0.01) reduced. Downregulation of NEU3 resulted in a decrease in NEU1 (p < 0.01) and NEU4 (p < 0.05) mRNA expression. In the NEU4 knockdown samples, NEU1 mRNA expression was significantly reduced (p < 0.05). Loss of function of NEU2 had no significant impact on mRNA expression of the other NEUs. Importantly, there is very little sequence homology between each NEU [23], and our BLAST results indicated that each of our siRNAs was specific to each NEU, thus, it is not likely that these results are due to off-target effects of our siRNAs. The results in DENV2 infected cells (Figure 5B) varied from the mock infected samples supporting our previous observations that NEU1-4 mRNA expression was altered upon DENV2 infection (Figure 4). In our NEU1 knockdown, we saw significant elevations in mRNA expression of both NEU2 (p < 0.001) and NEU4 (p <0.0001). In the NEU2 knockdown, NEU4 mRNA expression was reduced (p < 0.01). There were no changes to mRNA expression of other NEUs upon knockdown of NEU3. Finally, loss of function of NEU4 in DENV2 infected cells resulted in significant knockdown (p > 0.0001) of NEU1-3. Overall, our results indicated that only NEU4 seems to compensate for loss of function of NEU1, and, otherwise, expression of each NEU displayed some inter-dependence on expression of other NEUs. This suggested that NEUs or their downstream products may act as transcriptional regulators of one another.

**Figure 5.**
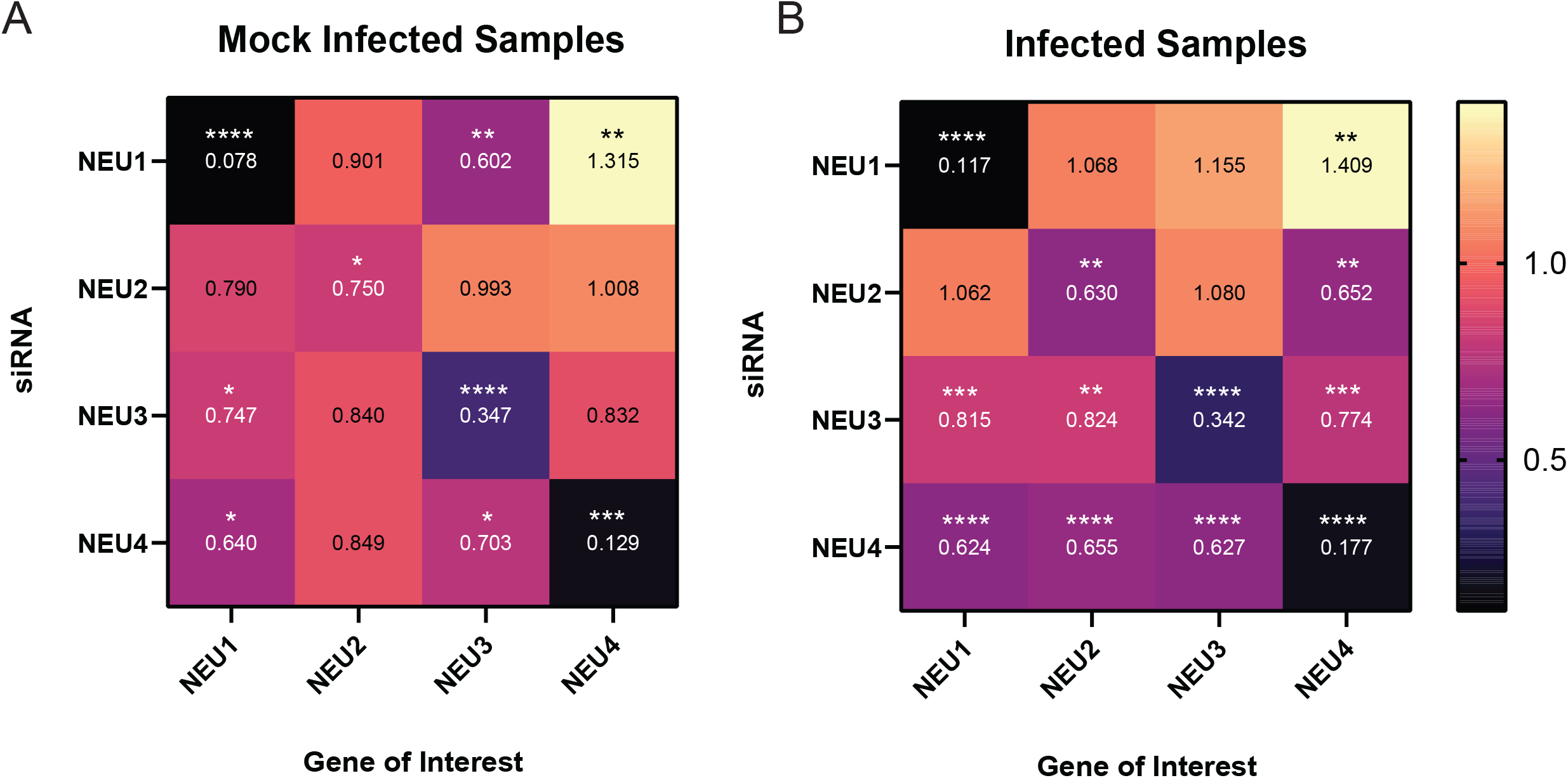
loss of function of NEU1-4 impacts mRNA expression of other NEUs. Huh7 cells were transfected with siRNAs targeting NEU1-4 and an IRR control siRNA, and either mock-infected (A) or DENV2-infected (B). At 24 hpi, cells were collected and analyzed via qRT-PCR to determine the relative mRNA expression of each NEU upon KD of other NEUs. Results were normalized to the RPLP0 housekeeping gene and compared to the IRR control (not shown) for each group. Results are displayed as a heatmap showing the fold change of each gene under each condition, written values inside boxes are average fold change values. Stars represent statistical significance in fold change compared to the corresponding IRR control. (A-B: One-way ANOVA with Dunnett’s multiple comparison’s test: * = p≤0.05, ** = p≤0.01, ***= p≤0.001, **** = p≤0.0001.)

## Discussion

Viruses such as DENVs are reliant upon and hijack host metabolic pathways to fulfill their replicative needs [6,24–29]. Dysregulation of host metabolic pathways leads to some of the pathogenesis seen in DENV infection, as was shown in studies that discovered NS1-mediated upregulation of sialidase activity was linked with endothelial hyperpermeability and vascular leakage [9,10]. However, a functional role for the increased sialidase activity remained unexplored. In this study, we sought to further characterize the relationship between DENV and the human sialidase enzymes. We have demonstrated that sialidase activity is vital for several steps in the DENV2 lifecycle. Specifically, we have shown that loss of function of NEU1-3 attenuates viral egress and to a lesser extent viral genome replication. In addition, we have now demonstrated that NEU4, previously unstudied in DENV infection, appears to not only be the most consequential sialidase to the DENV2 lifecycle, but that its expression is also necessary for the expression of NEU1-3. We also observed a periodicity to NEU3 expression during infection that seems to coincide with the start of a new replication cycle and may be NS1-mediated.

Recent studies have highlighted there is much ambiguity in the functional roles of NEUs. Each NEU retains some activity towards SIA residues beyond their preferred substrates [11,13,30,31], and some NEUs have been shown to be capable of translocating and acting in different subcellular compartments beyond their primary locations [21,32–34]. Moreover, the regulatory elements that govern NEUs remain unknown [35]. This level of ambiguity presented a challenge to determining the mechanism by which desialylation impacts DENV2 replication and viral release from cells. Thus, we chose to limit our study to determining which parts of the viral lifecycle were inhibited by loss of function of sialidase activity.

Several other possibilities that have emerged to explain the influence of sialidases on the DENV2 lifecycle. Desialylation of low density lipoproteins (LDLs) by NEU1 and NEU3 has been shown to increase cellular uptake of LDLs and increase intracellular cholesterol, possibly through the removal of steric hindrance on apo B-100, a ligand for the LDL receptor [36–39]. During DENV infection, LDL uptake increases at early timepoints of infection [40], thus, upregulation of NEUs may augment LDL uptake, providing DENVs with some of the lipid precursors necessary for membrane restructuring that drive formation of replication complexes. Another possibility is that intracellular trafficking along the endocytic and lysosomal pathways are, in part, modulated by the sialylation state of glycoconjugates within endocytic vesicles [41,42]. Importantly, DENVs are critically dependent upon endosomes to enter cells, have been shown to traffic between intercellular compartments using the endolysosomal system, and viral replication complexes have been found in components of the autophagolysosomal system [43–47]. Alternately, sialidases may act on the conserved glycosylation site at N154 (in most flaviviruses, N153) in the viral envelope (E) protein that has been proposed to shield the fusion loop preventing premature fusion [48–50]. Thus, sialidase activity in the late endosome may partially drive the fusion event necessary to release the viral RNA into the cytoplasm. Taken together, this suggests a model by which cellular sialidase activity supports the events necessary for viral uncoating, the membrane remodeling necessary to support viral replication, and intracellular trafficking of virus particles within infected cells. Further studies are necessary to parse these or other potential mechanisms by which NEUs influence the DENV2 lifecycle.

In summary, we provided the first known evidence of a transcriptional co-dependence between the human sialidase enzymes. Importantly, although previous studies established that the DENV NS1 protein modulated NEU1-3 activity, our present study highlights that manipulation of these enzymes, and thus cellular glycosylation, benefits DENVs. Future studies using NEU knockout cell lines, coupled with LC-MS/MS analysis of cellular glycoconjugates may help delineate which glycoconjugates are desialylated during DENV2 infection shedding light on the mechanism by which desialyation benefits DENV2 replication.

## Materials and Methods

### Cell Lines and Viruses

The cell lines used were the following: Clone 15(ATCC CCL-10) of the Baby Hamster Kidney Clone 21 cells (BHK-21), human hepatoma cells (Huh7) (unknown sex, From Dr. Charles Rice) [51], African Green Monkey kidney epithelial (Vero) cells (ATCC CRL-1586), and C6/36 cells (ATCC CRL-1660, larva, unknown sex). Huh7 and Vero cells were maintained in Dulbecco’s Modified Eagle Medium (DMEM) (Gibco, LifeTech), while BHK-21 and C6/36 cells were maintained in Minimum Essential Media (MEM) (Gibco, LifeTech). All media was supplemented with 10% heat-inactivated fetal bovine serum (FBS) (Atlas Biologicals), 2 mM nonessential amino acids (HyClone), 2 mM L-glutamine (Hyclone), and C6/36 media was also supplemented with 25 mM HEPES buffer. Huh7, BHK-21, and Vero cells were maintained at 37°C with 5% CO_2_, and C6/36 cells were maintained at 28°C with 5% CO_2_.

The virus strain used was: DENV2 (16681) [52]. DENV2 was passaged in C6/36 cells. Viral titers were obtained via plaque assay on BHK-21 cells [53]. Viral infections were performed at room temperature for 1 hr to allow for viral adherence. Afterwards, virus was removed, cells were washed with 1X PBS, and cell were overlaid with MEM or DMEM supplemented with 2% FBS, 2mM non-essential amino acids, and 2 mM L-glutamine. Cells were incubated at 37°C with 5% CO_2_ for indicated time periods.

### RNA extraction and qRT-PCR

Standard TRIzol extraction methods using either TRIzol or TRIzol LS (ThermoFisher) were used to extract RNA from cells and from viral supernatant. The Brilliant III Ultra-Fast SYBR® Green one-step qRT-PCR kit (Agilent) was used for all qRT-PCR reactions. The cycling parameters used were as follows: 20 mins at 50°C for reverse transcription, followed by 5 minutes at 95°C, and then 45 two-step cycles of 95°C for 5 secs and 60°C for 60 secs. A melt curve followed each cycle starting at 65°C and ending at 97°C. (Primer sequences are reported in Supplemental Table 1) To quantify DENV2 genome copies, *in vitro* transcribed viral RNA from a DENV2 cDNA subclone was used to generate a standard curve. All cellular RNA was normalized to Ribosomal Protein Lateral Stalk Subunit P0 (RPLP0) using the delta delta ct method as described in [6,54].

### siRNA transfection and confirmation of loss of function

Loss of function of NEU1-4 was conducted by transfecting cells with pooled siRNAs (Supplemental Table 1) (Dharmacon) or single siRNAs (Supplemental Table 1) (ThermoFisher) using RNAiMAX (Invitrogen) as described in [55]. Subsequently, cells were either infected with indicated viruses (described above), assessed for cytotoxicity of siRNA treatment (described below), or collected for confirmation of gene knockdown. Following indicated timeframes, viral supernatant and cells were collected. Viral supernatant was titrated via plaque assay. RNA from cells was analyzed via qRT-PCR to confirm knockdown of mRNA transcripts as described above. All siRNA treated samples were also compared to an irrelevant (IRR) siRNA control. Cytotoxicity of siRNA treatment was determined by replacing cell media with a 1:10 dilution of alamarBlue (ThermoFisher) in appropriate cell culture media and incubating for 1-2 hrs. A Victor 1420 Multilabel plate reader (Perkin Elmer) was used to measure fluorescence output with excitation at 560 nM and emission at 590 nM.

### Western Blot Analysis

Huh7 cells from IRR control and NEU1-4 KD samples were lysed in RIPA Buffer. Total protein in each sample was measured using the Pierce™ BCA Protein Assay Kit (ThermoFisher) according to manufacturer’s instructions. Equal total protein was loaded onto on a Criterion™ XT 4-12% Bis-Tris protein gel (Bio-Rad) and the was run for 2 hours at 100V. Protein was transferred at 4°C to nitrocellulose membrane for 2 hours at 50V. Following transfer, blots were blocked overnight at 4°C in a 5% milk solution in 1x PBS supplemented with 0.1% Tween 20. The primary antibodies used were a 1:500 dilution of the flavivirus group antigen antibody (D1-4G2-4-15, or 4G2) which binds domain II of the DENV E protein (mouse monoclonal antibody, BioVision) and 1:100 dilution of β-actin (rabbit polyclonal antibody, Invitrogen). A 1:3000 dilution of goat-anti-mouse IRDye 800CW, and goat-anti-rabbit IRDye 680RD (Li-Cor Biosciences) were used as secondary antibodies. Blots were imaged on a ChemiDoc MP Imaging System (Bio-Rad), and quantified in ImageJ utilizing area under the curve analysis.

### Immunofluorescence Assays

Huh7 cells were seeded in 12 well plates containing sterilized glass cover slips, and subjected to siRNA transfection and viral infection as described above. At 24 hpi, viral supernatants were collected and used to confirm reduction in viral release via plaque assay on BHK-21. Cells were then washed twice in 1x PBS, and fixed with a 4% paraformaldehyde solution for 20 minutes at room temperature, and then quenched with a 30 mM glycine/PBS (1x, pH 7.5) solution for 5 minutes. Following fixation, indicated samples were permeabilized in a 0.1% Triton X-100 (Sigma Aldrich), 1% Bovine Serum Albumin (BSA, Gold Biotechnology) in 1x PBS solution at room temperature, and then blocked with a 0.01% Triton X-100, 1% BSA in 1x PBS solution overnight at 4°C. Non-permeabilized samples were blocked with a 1% BSA in 1x PBS solution overnight at 4°C. The primary antibodies used were a 1:1000 dilution of 4G2 (mouse monoclonal antibody, described above), a 1:500 dilution of anti-Glut1 (rabbit monoclonal antibody, ThermoFisher) or a 1:500 dilution anti-Calnexin (rabbit polyclonal antibody, ThermoFisher). Glut1 and Calnexin were used as cellular markers for the plasma membrane and endoplasmic reticulum, respectively. For staining with neuraminidase enzymes, cells were fixed, permeabilized and blocked as described above. The primary antibodies used were a 1:1000 dilution of 4G2 (mouse or rabbit monoclonal, described above), and 1:50 dilutions of either anti-NEU1 (rabbit polyclonal, ThermoFisher), anti-NEU2 (mouse monoclonal, Santa Cruz Biotechnologies), anti-NEU3 (mouse, monoclonal, MBL international) or anti-NEU4 (rabbit polyclonal, Epigentek). The secondary antibodies were a 1:500 dilution of goat anti-mouse Alexa Fluor 647 (ThermoFisher) and 1:500 dilution of goat anti-rabbit Alexa Fluor 488 (ThermoFisher). Coverslips were counterstained with a 1:1000 dilution of the DAPI nuclear stain, and then fixed to slides with FluoroSave Reagent (Calbiochem). Slides were imaged using an Olympus inverted IX81 FV1000 laser scanning confocal microscope under the 100x oil objective using Fluoview FV10-AS2 4.2 software (Olympus). Images were processed with Volocity software version 6.3 (PerkinElmer).

### 3.4.7 Quantification and statistical analysis

Statistical details are noted in each figure and corresponding figure legend. Results are expressed as mean values with standard deviation. Statistical significance was determined using either a one-way Analysis of Variance (ANOVA) with Tukey’s or Dunnett’s multiple comparisons test using Prism software version 9.0 (GraphPad Software, La Jolla, California, USA).

## Acknowledgments

We would like to acknowledge and thank Elena Lian for providing critical evaluation of the data throughout this project.

## Author contributions

LAS and RP contributed to conceptualization and development of methodology for this manuscript. LAS, PP, and RP contributed to data acquisition and analysis. LAS and RP contributed to writing, reviewing and editing of the manuscript.

## Figure Legends

**Supplemental Figure 1 – Confirmation of knockdown of siRNA-mediated knockdown of sialidase mRNA levels**

(A-B): Huh7 cells were transfected with pooled siRNAs targeting NEU1-4 and the indicated controls followed by mock-infection (A) or DENV2-infection (B). At 24 hpi, cells were collected, processed, and relative mRNA expression was analyzed via qRT-PCR. Values were normalized to the RPLP0 housekeeping gene. (A-B: one-way ANOVA with Tukey’s multiple comparisons test: * = p≤0.05, ** = p≤0.01, ***= p≤0.001, **** = p≤0.0001.)

**Supplemental Figure 2 – DENV2 E protein accumulates in cells with reduced NEU1-4 expression**

(A-D) Huh7 cells were transfected with siRNAs targeting NEU1-4 and or non-targeting siRNA control (IRR), followed by infection with DENV2 (MOI = 0.3). At 24 hpi, cells were fixed in 4% paraformaldehyde, and permeabilized to determine whether NEU1-4 KD inhibits viral egress from cells. Primary antibodies used were mouse anti-4G2 (DENV2 E protein, A,D) or rabbit anti-4G2 (DENV2 E protein, B,C), rabbit anti-NEU1 (A), mouse anti-NEU2 (B), mouse anti-NEU3 (B), and rabbit anti-NEU4 (D). Cells were counterstained with goat anti-mouse AlexaFluor 488, goat anti-rabbit AlexaFluor 647, and DAPI nuclear stain and imaged on an Olympus inverted laser scanning confocal microscope and processed using Volocity software v6.3.

