## Supplemental Figures and Tables for "Human sialidase activity is vital for dengue virus serotype 2 infection"

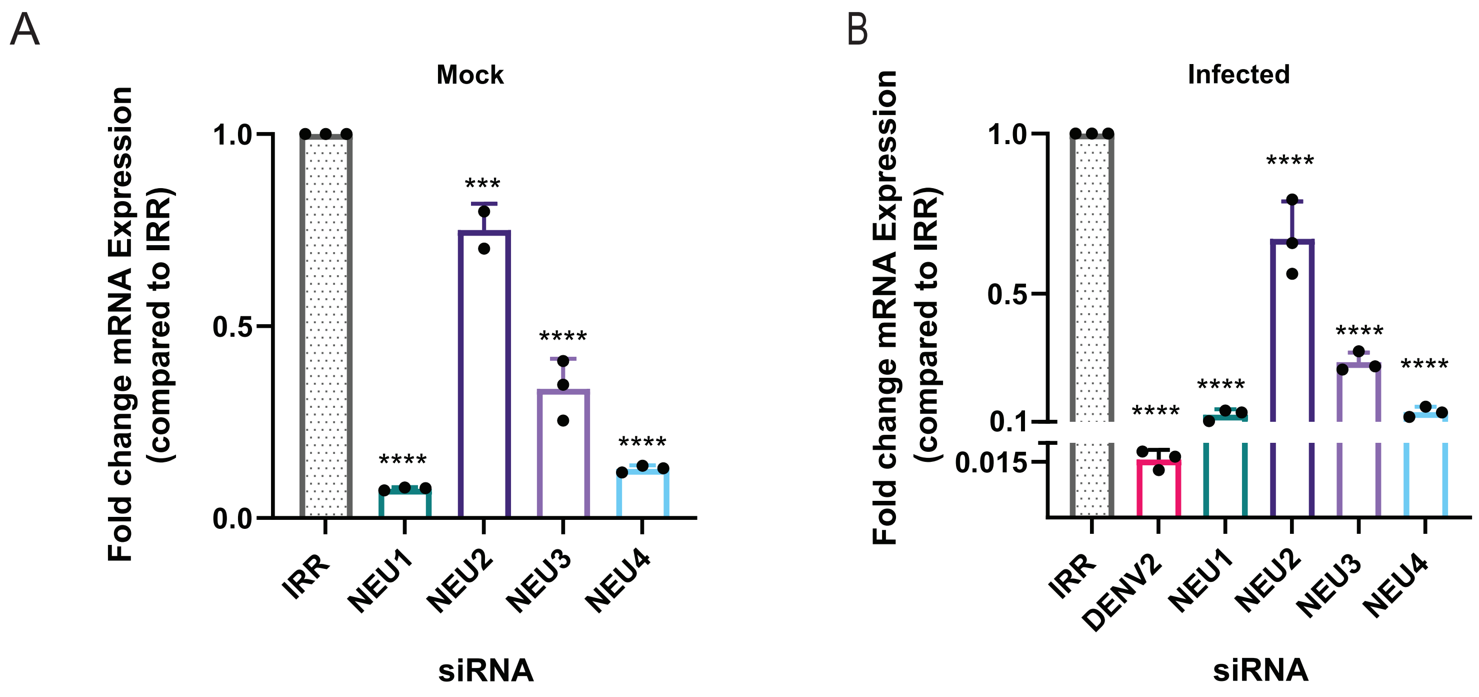

Supplemental Figure 1, St Clair LA, et al.

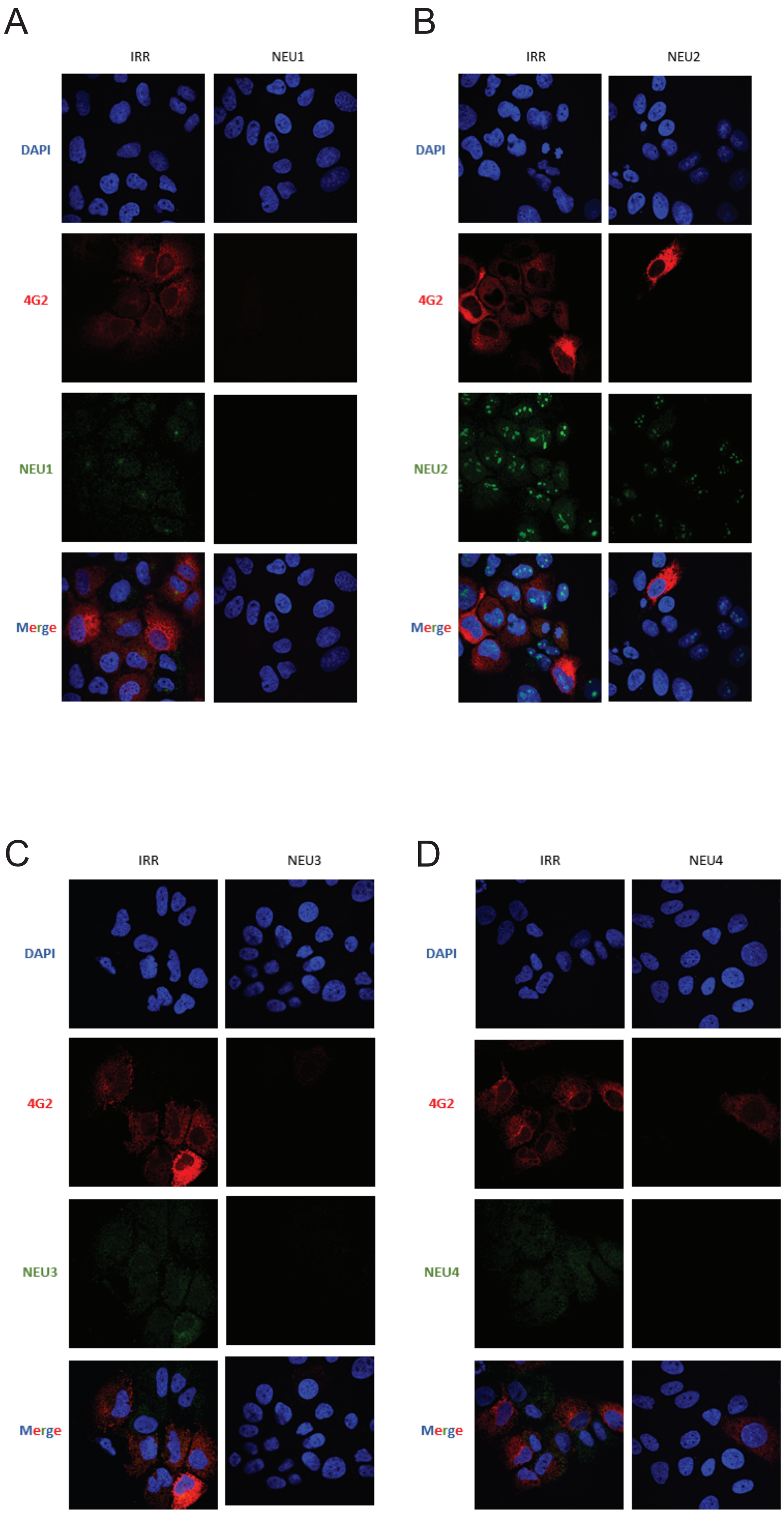

Supplemental Figure 2, St Clair LA, et al.

**Supplemental Table 1. siRNA reagents and primer sequences used in this study.**

| **siRNA reagents and sequences** | | | |
|---|---|---|---|
| **siRNA** | **Manufacturer** | **Product Identifier** | **Sequence(s)** |
| Irrelavant control siRNA (custom) | Horizon Discovery | Custom siRNA | GGACUUCCAGAAGAACAUCTT |
| DENV2 control siRNA (custom) | Horizon Discovery | Custom siRNA | CGGGAAAGACGAAGAGAUAUU |
| Neraminidase 1 (SMARTpool) | Horizon Discovery | M-011092-00 | GAAAGAGACAGUCCAGCUA |
| | | | CGGAAUCUCUCCCUGGAUA |
| | | | CGAUGGAGCUUCAGCAAUG |
| | | | AGUGAGCGAUGUUGAGACA |
| Neraminidase 2 (SMARTpool) | Horizon Discovery | M-012058-00 | GUACGAAGCCAAUGAUUAC |
| | | | CCAAUGACGGGCUUGAUUU |
| | | | CAAGAAGGAUGAGCACGCA |
| | | | GGCAAGUCACGGAGCAACA |
| Neraminidase 3 (SMARTpool) | Horizon Discovery | M-010641-01 | GAAGAUGACAGAGGGAUUA |
| | | | GAUCUACAGUGAUGACCUA |
| | | | GUGCAGAGGUCAUGGAAGA |
| | | | GAACCCAAGCCAAUUCAAA |
| Neraminidase 4 (SMARTpool) | Horizon Discovery | M-013263-00 | GAUGAGAUUUCCUUUUGUA |
| | | | GUGCAGAUCGCCACGGGAA |
| | | | GCUCGGCUACACAUGGGUA |
| | | | GGCCACGGGAUGACAGUUG |
| NEU1 silencer siRNA | ThermoFisher | AM16708 / assay ID 8573 | AUUUCUUUUCUACUCCCUUU |
| NEU2 silencer siRNA | ThermoFisher | AM16708 / assay ID 45117 | UUACGAGGAGAUUGUCUUUC |
| NEU3 silencer siRNA | ThermoFisher | AM16708 / assay ID 135986 | CAGUUGGUACAGUGGGGGCC |
| NEU4 silencer siRNA | ThermoFisher | AM16708 / assay ID 36069 | UCUUUAGGAAGGGGAGCAGC |
| **Primer Sequences** | | | |
| **Target** | Abbreviation | **Forward** | **Reverse** |
| Ribosomal protein lateral stalk subunit P0 | RPLP0 | AGATGCAGCAGATCCGCAT | GGATGGCCTTGCGCA |
| Hexokinase II | Hexokinase | ATCCCTGAGGACATCATGCGA | CTTATCCATGAAGTTAGCCAGGCA |
| Neuraminidase 1 | NEU1 | GTCAGCCCAAGCAGGAAAATG | CGGGCATTGATGACGACTGA |
| Neuraminidase 2 | NEU2 | GCGCAGAGGAGACTACGAC | GTCATACAAGGGGCATGGGTT |
| Neuramindase 3 | NEU3 | GACCATGAACCCCTGTCCTG | AGCATTCCTGCCTGACACAA |
| Neuraminidase 4 | NEU4 | ACCGCCGAGAGTGTTTTGG | CGTGGTCATCGCTGTAGAAGG |
